# A frailty index for UK Biobank participants

**DOI:** 10.1101/233692

**Authors:** Dylan M. Williams, Juulia Jylhävä, Nancy L. Pedersen, Sara Hägg

## Abstract

**Background:** Frailty indices (FIs) measure variation in health between aging individuals. Researching FIs in resources with large-scale genetic and phenotypic data will provide insights into the causes and consequences of frailty. Thus, we aimed to develop an FI using UK Biobank data, a cohort study of 500,000 middle-aged and older adults.

**Methods:** An FI was calculated using 49 self-reported questionnaire items on traits covering health, presence of diseases and disabilities, and mental wellbeing, according to standard protocol. We used multiple imputation to derive FI values for the entire eligible sample in the presence of missing item data (N =500,336). To validate the measure, we assessed associations of the FI with age, sex, and risk of all-cause mortality (follow-up ≤ 9.7 years) using linear and Cox proportional hazards regression models.

**Results:** Mean FI in the cohort was 0.125 (standard deviation = 0.075), and there was a curvilinear trend towards higher values in older participants. FI values were also marginally higher on average in women than men. In survival models, 10% higher baseline frailty (i.e. a 0.1 FI increment) was associated with higher risk of death (hazard ratio (HR) = 1.65; 95% confidence interval: 1.62, 1.68). Associations were stronger in younger participants than in old, and in men compared to women (HRs: 1.72 vs. 1.56, respectively).

**Conclusions:** The FI is a valid measure of frailty in UK Biobank. The cohort’s data are open-access for researchers to use, and we provide script for deriving this tool to facilitate future studies on frailty.

## Introduction

Frailty has been defined as the variation in health between individuals as they age.(1) A common approach for measuring frailty is the calculation of a frailty index (FI), based on the proportion of measured health deficits that individuals accrue with age.(2) There are well-documented properties of the FI, such as an established association of higher FI scores with increased all-cause mortality risk, and with risk of age-related diseases and clinical endpoints.(3-6) FIs also have demonstrable consistency in these respects despite being calculated across different research resources, with varying numbers and types of measures used for FI composition.(7)

Aside from suggested clinical applications, FIs have also been considered for studying the biological basis of ageing.(8) Further research using FI as a proxy for biological age could shed light on genetic and environmental determinants of age-driven health deterioration, and the etiologies of many age-related diseases.

To address this opportunity, we aimed to develop and validate an FI using baseline assessment data from UK Biobank (UKB): a cohort study of approximately half a million adult participants with data available openly for researchers.(9) In this article, we describe the construction of an FI using UKB data, its characteristics, and test whether the measure is associated prospectively with risk of all-cause mortality. We provide the script for the FI derivation in UKB as a tool to facilitate future research on frailty using this study resource.

## Methods

### Participants

UKB is a multi-centre cohort study with 502,631 participants, aged 40 to 69 years, enrolled at 22 assessment sites in England, Scotland and Wales between 2006 and 2010.(9) All participants undertook a baseline assessment, which involved physical measures, biological sample collection for biobanking, and verbal interviews and touch-screen questionnaires that addressed numerous traits (including demographics, socioeconomics, lifestyle, environmental exposures, and health factors and medical history). The UKB sample has been followed up for mortality and subsequent development of diseases via linkage to information held in national death registers and by hospital episode statistics. UKB data are open-access for all researchers with an approved research proposal, and the entire data catalogue can be browsed via the cohort’s website: http://www.ukbiobank.ac.uk/.

In this study, we used questionnaire and interview data from the cohort’s baseline assessment to derive a standard FI. We also used mortality data and dates of death from death certificates held by the National Health Service (NHS) Information Centre (for participants in England and Wales) and NHS Central Register (for those in Scotland). Participants were excluded from the sample if missing data for 10 or more items (over 20%) that were used to calculate the FI. We also excluded three participants who had death certificate dates that preceded dates of attendance at the UKB baseline assessment. After exclusions, the eligible analysis sample was 500,336. The sample derivation is depicted in supplemental figure 1.

### Frailty index construction

FI values were calculated for participants according to a standard protocol.(2) Deficits are based on indicators of ill health across a variety of physiological and mental domains, and can include symptoms, diagnosed diseases, and disabilities. The FI can combine continuous variables with categorical or binary variables by assigning values for each trait between zero and one according to the severity of the deficit (zero meaning a deficit is absent, and one meaning the deficit is at its most severe). An FI value is then composed as the sum of deficits accrued by an individual divided by the total number of deficits composing the FI, e.g. an individual with 10 deficits from a total of 50 items, would have an FI value of 0.2 (10/50).

Criteria for inclusion of variables as items were as follows, adapted from (2):

- Traits should be health deficits, i.e. *not* lifestyle characteristics or behaviors related to disease or mortality risk, such as smoking.
- Risk of the deficit should be higher with increasing age.
- Traits should not be very rare, i.e. a deficit should not have a prevalence within the sample below 1%.
- Traits should cover a range of physiological areas, e.g. physiological aspects, physical function and mental wellbeing.
- Traits should not be ubiquitous in the (baseline) sample, e.g. long-sightedness being almost universal in midlife.
- Data on the traits should be recorded for at least 80% of individuals.

Following a search of all self-reported baseline data available against these criteria, 49 variables were identified for inclusion as FI items. The items and their coding are detailed in supplemental table 1. To test whether any items should be excluded from the FI due to lack of independence with respect to multiple items, we examined item pair correlations by Pearson and Spearman’s rank correlation coefficients (for pairs including binary / categorical items, respectively). All between-item correlations were moderate or low (all *r* or rho <0.43), so no items were excluded on this basis.

In addition to variables used to code FI items, we also used information on sex and ethnicity for covariates in statistical models. Responses regarding ethnicity were combined into six major categories: ‘white’, ‘black or black British’, ‘Asian or Asian British’, ‘Mixed’, ‘Chinese’ or ‘Other’, as per the cohort’s groupings.

### Missing data

Missing data would have precluded analyses for 19.6 percent (N=96,197) of otherwise eligible participants in the cohort, so we imputed data and conducted analyses on the entire sample. Data were imputed for individuals without values for one to nine of the FI items and/or data on ethnicity using multiple imputation (MI) by chained equations.(10) More information on the MI modelling is provided in the supplementary material. Main analyses were based on the imputed data, and the models were repeated in complete-case data for comparison.

### Statistical analysis

To validate the FI, we plotted its distribution and examined its associations with age, sex and all-cause mortality using regression models. In all models, the FI variable was entered without transformation. Non-linear trends of higher FI values with increasing age have been observed previously in both cross-sectional and longitudinal samples.(11) We therefore tested for a non-linear association of the FI (entered as the dependent variable) with age using fractional polynomial statistics and related plots.(12) To avoid the inclusion of all polynomial terms in the imputation of the full sample, the fractional polynomial regression was conducted after MI in the first imputed dataset alone, and repeated in other imputed sets and the complete-case data to check consistency.

For modeling of the association of FI with mortality risk, attained age was used as the time scale to account for variable ages at entry (38 to 73 years), staggered baseline assessments between 2006 and 2010, and varying lengths of follow-up time thereafter (≤ 9.7 years). Risk estimates based on this scale are implicitly adjusted for differences in age at baseline. Calculation of attained age was based on date of birth as the origin point, date of baseline attendance as the entry point, and exit at date of death or censorship on 30^th^ November 2015 (the latest date of complete coverage across death registers used for UKB data at the time of analysis). We assigned the 15^th^ day of each month as the day of birth for all participants, since we had data on month and year of births only. Association of baseline FI variation with risk of all-cause mortality was assessed by Cox proportional hazards regression. We conducted three models: 1) with the FI as a single independent variable; 2) with additional adjustment for sex; 3) with both sex and ethnicity as covariates. We scaled the models so that hazard ratios are expressed per 0.1 higher frailty (equating to a 10% increase). We plotted Kaplan-Meier survival curves by five groups of FI values at 0.1 increments across the observed FI distribution (<0.1, 0.1 to < 0.2, 0.2 to < 0.3, 0.3 to < 0.4, and ≥ 0.4). We examined for violation of the proportional hazards assumption for Cox regression models plotting ‘log-log’ values for survival curves over analysis time by the five FI categories, and of the scaled Schoenfeld residuals from Cox models regressed on analysis time.

### Secondary analyses

The suitability of the FI for measuring biological age on individuals during young adulthood and middle-age (when age-related clinical manifestations are more scarce) has been questioned.(13) Thus, we investigated the association of the FI with mortality stratified by age group at which participants were recruited. We categorized age at baseline into the following groups, in which at least 1000 deaths per group had arisen by the point of censorship: <50, 50 to <60, 60 to <65, and ≥65.

We tested for an interaction of the FI-mortality association by sex in the whole sample, and then conducted an additional analysis for survival models stratified by sex. We also tested for an interaction by ethnic group of participants.

All analyses were conducted in Stata version 15.0 (StataCorp LLC, College Station, Texas, USA).

### Ethics

All individuals in the study sample gave written informed consent to participate in UKB and for data to be used in future research. Ethical approval for this study is covered by the general ethics review for UKB, conducted by the North West Haydock Research Ethics Committee of the UK’s Health Research Authority (Reference 16/NW/0274, 13^th^ May 2016).

## Results

Characteristics of participants in the full analysis sample after MI (and stratified by age categories) are shown in table 1. Supplemental table 2 shows these characteristics by groups with and without missing data. There were discernible differences in the proportions of female participants, distributions of ethnic groups, and proportions that died – indicating the benefit of performing MI on the eligible sample. FI values for the complete-case sample were also expected to be lower than corresponding values for those with missing data.

**Table 1:**
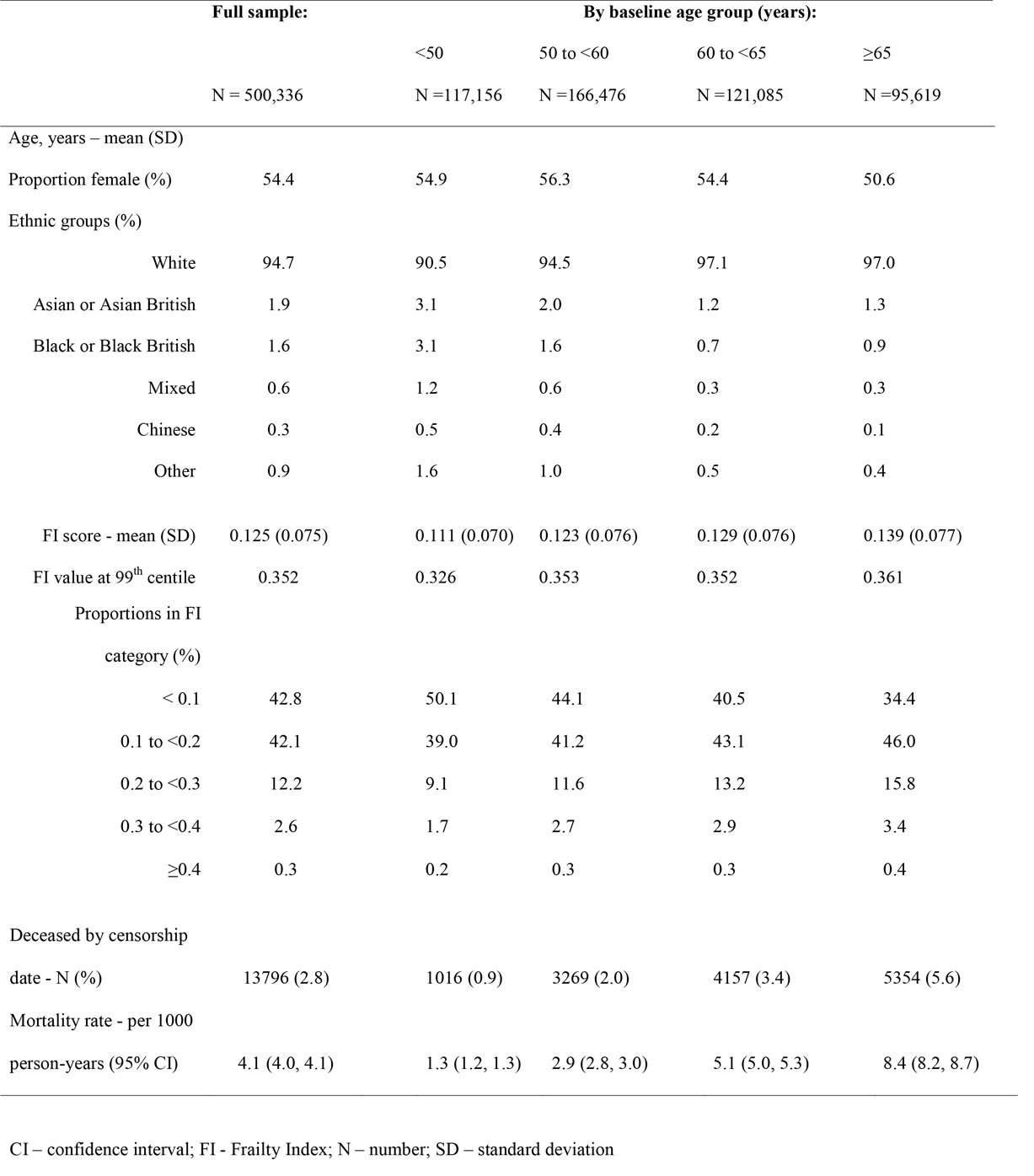
Characteristics of UK Biobank cohort participants, in the full analytical sample, and stratified by age groups

The distributions of the FI in men and women are depicted in supplemental figure 2. The difference in FI values between sexes was not pronounced, though women had higher values across the right tail of the distribution than men. Univariable regression of the FI on sex indicated that, on average, FI values were 0.78% higher in women than men (95% confidence interval (CI): 0.74, 0.82%). The 99^th^ centile was 0.352, and the highest FI value in the full sample (0.600) was within the maximum empirical limit for FI values (under 0.7) that has been suggested consistently by past studies.(14)

The use of fractional polynomials suggested a slight curvilinear relationship between FI values and age at the baseline assessment (figure 1). The best fitting regression equation of 44 combinations tested was to model age with two terms raised to powers −2 and 3. There was strong statistical evidence for this fit differing from both a linear model (P<0.001) or a model including an additional quadratic term (P=0.001). Results were almost identical in imputed and complete-case data.

**Figure 1:**
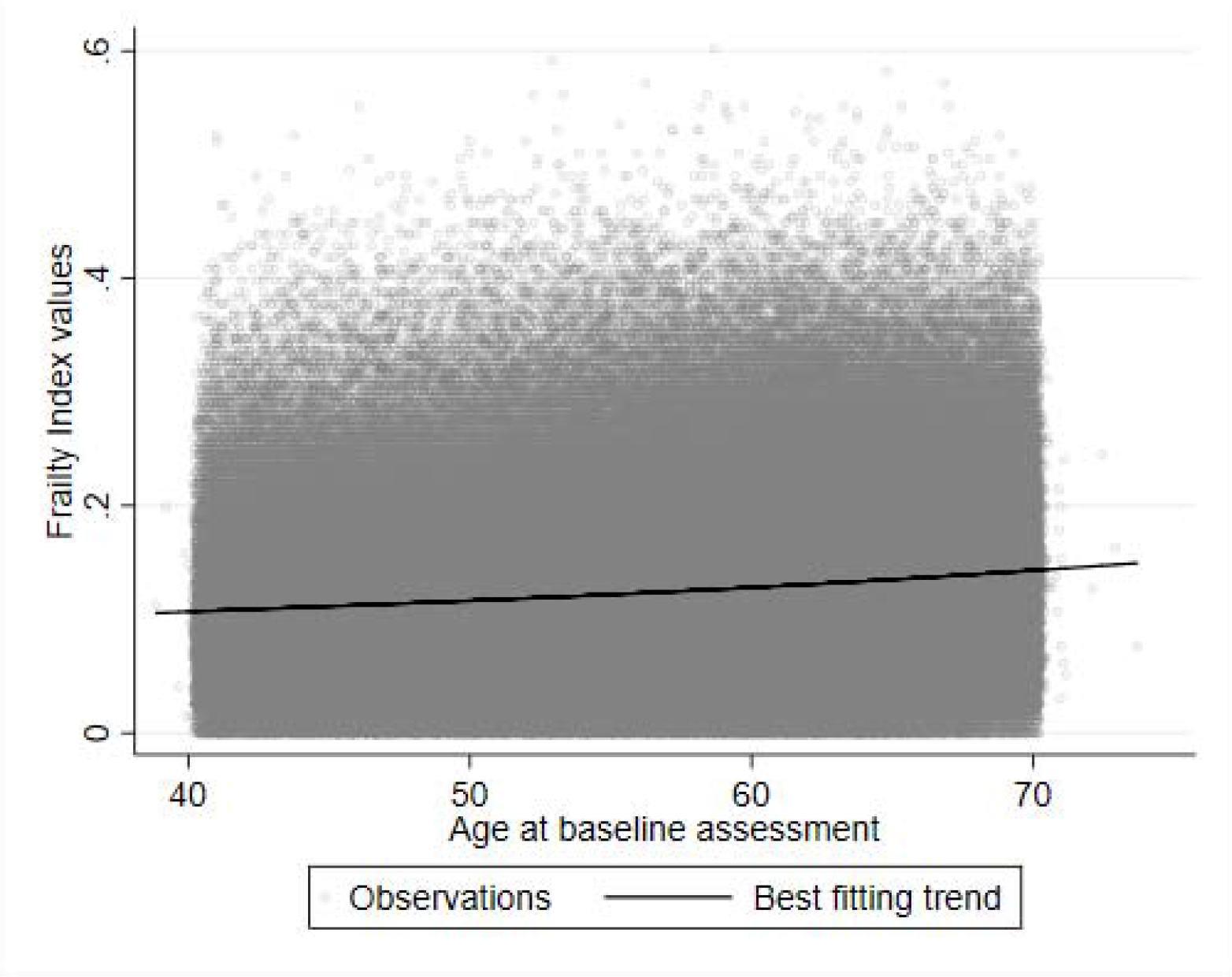
Distribution of FI values by age at baseline (N=500,336)

The results of Cox proportional hazards regression models for the association of baseline FI values with survival after a maximum of 9.7 years are shown in table 2. Higher frailty values were strongly associated with mortality, and the magnitudes of association increased slightly with adjustment for sex and ethnicity (models 2 and 3). Scaled by 0.01 higher FI values, results for models 1 to 3 were 1.049 (95% CI: 1.047, 1.051), 1.051 (1.049, 1.053) and 1.051 (1.049, 1.053), respectively. In survival models stratified by ages at baseline, higher FI values were most strongly associated with mortality in younger age groups. There were no obvious departures from proportional hazards in Cox models. Results from the complete-case sample did not differ notably from the imputed sample findings, e.g. the hazard ratio from model 1 (adjusted for age only) was 1.62 (95% CI: 1.59, 1.66).

**Table 2:** Hazard ratios for mortality according to baseline FI, after ≤ 9.7 years of follow-up in UK Biobank

|  | Model 1 |  |  | Model 2 |  | Model 3 |  |
| --- | --- | --- | --- | --- | --- | --- | --- |
|  | Deaths (N) | HR | 95% CI | HR | 95% CI | HR | 95% CI |
| Full sample (N=500,336): | 13,796 | 1.61 | (1.58, 1.64) | 1.65 | (1.62, 1.68) | 1.65 | (1.62, 1.68) |
| Females only (N=272,301): | 5422 | 1.55 | (1.50, 1.60) | - | - | 1.56 | (1.51, 1.60) |
| Males only (N=228,035): | 8374 | 1.71 | (1.67, 1.76) | - | - | 1.72 | (1.68, 1.76) |
| Age group at baseline: |  |  |  |  |  |  |  |
| <50 (N =117,156) | 1,016 | 1.84 | (1.72, 1.98) | 1.86 | (1.74, 2.00) | 1.87 | (1.74, 2.00) |
| 50 to <60 (N =166,476) | 3,269 | 1.73 | (1.66, 1.79) | 1.76 | (1.70, 1.82) | 1.77 | (1.70, 1.83) |
| 60 to <65 (N =121,085) | 4,157 | 1.58 | (1.52, 1.63) | 1.60 | (1.55, 1.66) | 1.60 | (1.55, 1.66) |
| $\geq 65$ (N =95,619) | 5,354 | 1.54 | (1.49, 1.58) | 1.58 | (1.54, 1.63) | 1.59 | (1.54, 1.64) |

Results are expressed per 0.1 increments on the FI scale (10% higher frailty)
CI – confidence interval; FI – Frailty index; HR – Hazard ratio.
Model 1 – FI entered as the sole independent variable
Model 2 – sex included as an additional covariate (not conducted for sex-stratified samples)
Model 3 – as model 2, plus adjustment for ethnicity
Age is adjusted for in all models via the ‘attained age’ scale used for survival analyses

Figure 2 depicts Kaplan-Meier survival curves (unadjusted for covariates) that illustrate the estimated probability of survival for individuals throughout follow-up, given their age and category of FI values at the baseline assessment. There was a gradient of decreased life expectancy in those in the higher FI categories compared to lower categories, which widened with increasing age at baseline.

**Figure 2:**
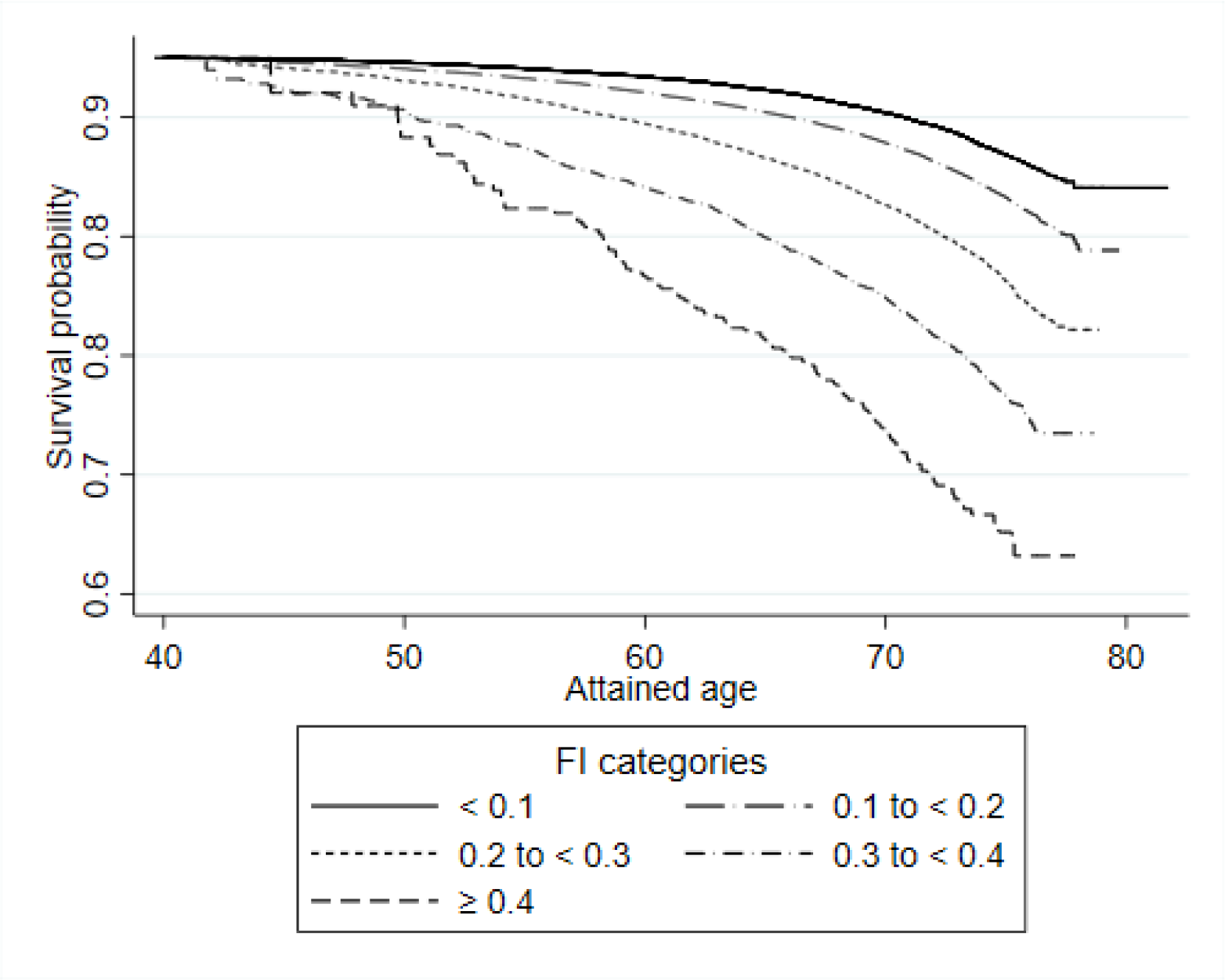
Survival curves according to baseline age and FI categories, after ≤ 9.7 years of follow-up (N=500,336) The age scale shows risk of mortality within the follow-up period according to the age at which individuals entered the cohort and their baseline FI category. For example, the average probability of survival after 9.7 years for a 70 year old in the lowest FI category was approximately 90%.

There was evidence that the FI-mortality association is modified by sex (*P* for interaction test <0.001), and in stratified models, the association was stronger in men than in women. There was weaker evidence that the FI-mortality association is modified by ethnicity (*P* = 0.10), so we did not stratify results by ethnic group. However, there may have been limited power to detect such an interaction given the few deaths currently recorded for non-white ethnic groups, which are small sub-groups of the sample.

## Discussion

We have designed and validated a standard measure of frailty for half a million participants of the UKB cohort. The FI measure displayed all the expected characteristics in terms of its observed distribution, differences by age and sex, and its association with all-cause mortality risk – not only in old participants but also in those who were recruited in middle-age. These factors illustrate the utility of the FI for future studies of frailty using UKB data.

There are many similarities of past findings with results from UKB. The high degree of consistency is expected, considering the insensitivity of FIs to the number and exact choices of items used to derive the composites(15) and observed similarities across geographical regions and demographic groups.(16) For instance, the difference in the FI between the sexes is commonly observed,(17) mirroring the paradox that women live longer than men on average despite suffering a higher burden of co-morbidity.

A meta-analysis of associations of FIs with survival has also highlighted the similarity of the measure as a surrogate for mortality risk across research settings.(7) The combined estimates yielded hazard ratios centred on 1.04 (95% CI: 1.03, 1.04) and 1.28 (1.26, 1.31) per 0.01 and 0.1 higher FI values, respectively. These are more modest than the associations observed in UKB, with equivalently scaled hazards being 1.05 and 1.65 in fully adjusted models. One explanation for this disparity could be the inclusion of middle-aged participants in UKB, since FI values were more strongly associated with mortality risk in younger age groups. Although the meta-analysis authors reported no evidence for modification of FI-mortality associations by age across all pooled studies, it is noteworthy that the strongest associations among individual samples were those that included middle-aged individuals in addition to older participants.(7) This phenomenon has been observed previously, but not with the statistical precision that the UKB sample provides when stratified into age groups.(18, 19) The finding may imply that FIs recorded in middle-age can proxy survival more effectively than in older individuals, perhaps indicating the sensitivity of a measure during a period of the life course when clinically manifested chronic disease burden is generally low and only small proportions of severely ill individuals are susceptible to death. This finding should be explored further in other large datasets.

There are several strengths to this research. Deriving the FI in a cohort of this size will enable new inferences on frailty, especially as the UKB data is enriched – for example, coverage of all participants with genome-wide and exome sequencing data for genetic association studies.(20) Another advantage is a large number of questionnaire items used to construct the FI, covering a wide range of health domains. The use of MI to impute missing data allowed almost the entirety of the cohort’s sample to be leveraged in this analysis. MI has been used very infrequently in frailty research,(21) despite it being the gold-standard method for handling missing data in observational studies (improving the precision of estimates, mitigating bias from missing values, and with inferential models reflecting the uncertainty of imputing incomplete information on participants).(10) It may be preferable to a more commonly used approach – to create an FI where the sum of deficits is divided by the number of non-missing item data on each individual, i.e. with varying denominators used to calculate FI values between individuals. Models based on this form of FI would falsely assume that frailty has been measured with no missing data, and so there is more prospect of uncontrolled bias and overly-precise inference.

Some limitations should also be highlighted. Despite being designed as a population-representative cohort, UKB recruitment was influenced by selection bias: participants were healthier in several respects than expected given population averages of lifestyle or health traits (the ‘healthy volunteer effect’).(22) However, the mean and 99th centile of the FI for those aged 65 or older in UKB (0.139 and 0.361, respectively) were not substantially different to corresponding values derived from health record data on the general population of the UK (0.14 and 0.49), considering this other sample included individuals aged up to 95 years.(6) Moreover, the main consequence of this selection bias is to limit the use of UKB data for calculating disease prevalence or incidence rates, but not etiological studies where sufficient variations in exposures and outcomes exist (discussed at length in (22)). With high observed variability, the FI data in UKB therefore still appears to be valuable for research into the etiology of frailty. Conversely, uses of the data for the development or calibration of the FI as a prediction tool may be inappropriate – at least without weighting results for differences in disease or mortality incidence expected in a target population. A second limitation is that, at present, the FI is derivable on the whole cohort only using data from the baseline assessment. However, sizeable sub-samples of the cohort have repeat data measured, allowing for study of frailty changes in these individuals, and further follow-ups of the cohort are planned, meaning that longitudinal frailty assessment should become possible for most of the full cohort. A third constraint of the FI that we developed is the use of only self-reported questionnaire data. Objectively measured traits can also be enrolled as FI items.(23) UKB has a number of relevant clinical measures in this respect, plus assay data on multiple biomarkers being produced. Once available, future studies could enrol these new data into a more comprehensive FI (or using more advanced modelling, such as principal component analysis) to reconcile variation in clinically manifested traits and biological measures together.

In conclusion, UKB data provide a promising avenue of research for understanding the causes and consequences of frailty. In designing and validating a standard FI in this resource, and providing the information necessary for other research groups to derive this index, our hope is to facilitate use of the tool and hasten research into frailty. As UKB continually improves with new genetic and phenotypic data, these opportunities will grow considerably and should be embraced by gerontologists.

## Acknowledgments

We are grateful to Hannah Bower, Paul Lambert and Henric Winell (Department of Medical Epidemiology and Biostatistics, Karolinska Institutet) for their advice on the statistical methods used in this research.

This research has been conducted using the UK Biobank resource, as part of the registered project 22224.

## Abbreviations

FI: Frailty Index
NHS: National Health Service
UKB: UK Biobank

